# SpaceExpander: Automated Drafting and Evaluation of Markush Claims for Chemical Space Expansion

**DOI:** 10.64898/2026.04.09.716825

**Authors:** Rui Wu, Liyun Mao, Yanyan Diao, Honglin Li

## Abstract

Drafting Markush claims for chemical patents remains difficult because manual claim writing is slow, error prone, and often fails to capture related chemical space in a systematic manner. We developed SpaceExpander, a computational method that converts disclosed compounds into generalized Markush claims by extracting core scaffolds, defining variable positions, decomposing complex substituents, and expanding substituent space through fragment matching. We evaluated the method on 24 publicly available chemical patents and compared its performance with IntelliPatent. SpaceExpander achieved a mean atom level scaffold accuracy of 0.92 and exactly recovered the reference scaffold in 19 of 24 patents. By contrast, IntelliPatent could process only 2 patents from the same set, indicating more limited applicability to structurally diverse cases. We further examined practical claim coverage in a case study based on the Osimertinib patent. Using representative disclosed compounds as input, SpaceExpander drafted a Markush claim that covered 5 of 7 additional approved third-generation EGFR inhibitors beyond Osimertinib. These results show that SpaceExpander is a validated method for automated Markush claim drafting and chemical space expansion.

## 1. Introduction

Patents safeguard the structural novelty of core compounds in the chemical and pharmaceutical industries. The extent of protection, including potential derivatives with similar effects, is ultimately defined by the claim language [1-3]. The claims section defines the legal scope of protection, outlining the distinguishing features of the invention and restricting unauthorized exploitation [4-8].

One common strategy involves starting with a core compound and expanding coverage using Markush structures, which allow a single claim to encompass multiple chemical variants [9-13]. Strengthening pharmaceutical Markush patents is key to encouraging innovation, but the scope must match the actual contribution of patentee to avoid overreach and ensure fair protection. Drafting such claims is complex, relying on expert knowledge to describe diverse analogs, with manual processes prone to error and information leakage [14-16].

Existing tools primarily focus on interpreting and querying Markush structures, such as iMarVis supports visualization and nested R-group handling [17,18], SMIRKS-based algorithms enable structure mapping [19]; and the Periscope system enables generation, visualization, and searching of Markush structures [20]. IntelliPatent automates the extraction of Markush scaffolds and the generation of claims; however, it cannot process complex fragments not already present in its predefined library [21]. Similarly, the Markush Editor provided by ChemAxon offers comprehensive functionalities but is a commercial product that requires a paid license, restricting its accessibility and broader adoption [22, 23]. Nevertheless, several AI-powered patent drafting tools, such as Patentext, AutoPatent [24], and ClaimMaster, have emerged, yet none of these tools are capable of drafting pharmaceutical patents involving chemical structures or Markush claims.

In this manuscript, we present SpaceExpander, a computational framework for automated Markush claim drafting, as shown in Figure 1. Unlike existing methods that rely on limited predefined fragments, SpaceExpander combines scaffold extraction with recursive handling of complex R groups to support structurally diverse compounds. The framework offers three main methodological advances: 1) automated scaffold extraction, which identifies the shared core structure of the input compounds while preserving substituent positions and bond variations; 2) hierarchical R-group decomposition, which recursively resolves complex structural fragments into standardized textual descriptions; and 3) fragment-based expansion of substituent space, which broadens Markush representations by incorporating chemically related structural motifs. The practical utility of the framework was further supported by accurate scaffold recovery across diverse chemical patents and by expanded claim coverage in a real-world drug series, as illustrated in the Osimertinib case study.

**Fig. 1.**
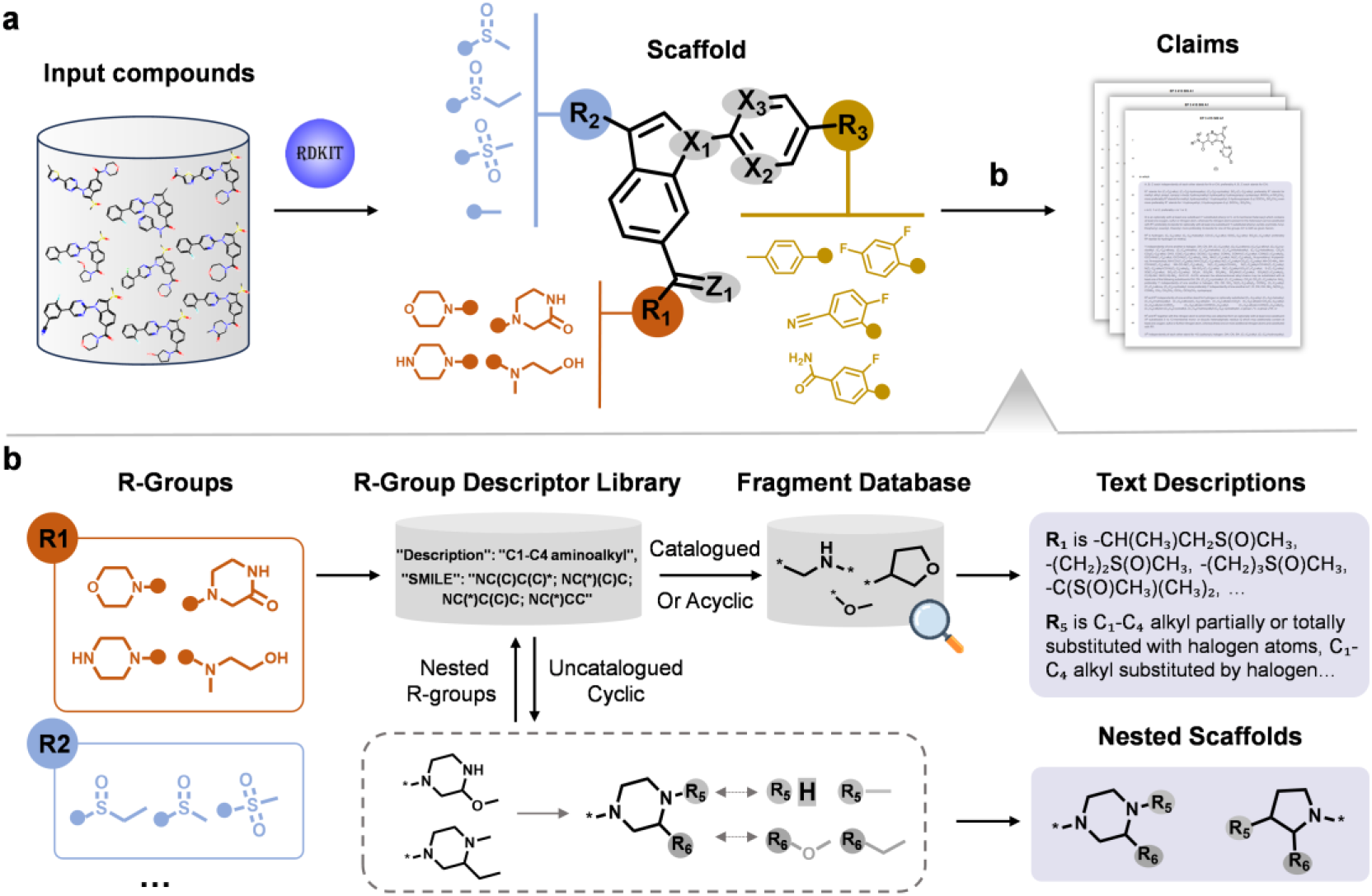
Overview of the study. (a) The system extracts a common scaffold containing placeholders (R, X, Z) from the input compounds, which are further processed in (b) to draft the final claims. (b) R-groups are matched against the R-group Descriptor Library. Catalogued or acyclic fragments are expanded via similarity-based retrieval from the Fragment Database and assigned predefined descriptions. For uncatalogued cyclic fragments, a Hierarchical Decomposition System recursively extracts nested scaffolds and nested R-groups, which are then returned to the Descriptor Library for further matching and expansion.

## 2. Methods

### 2.1 Extraction of optimal Markush scaffold

All input compounds are standardized into SMILES format and converted to Kekulé form to explicitly represent aromatic systems using alternating single and double bonds. Scaffold construction starts with identifying the shared core structure across compounds using the Maximum Common Substructure (MCS) algorithm.

The resulting scaffold is validated using two criteria: heavy atom count, to exclude undersized scaffolds (e.g., ≤3 atoms), and topological consistency, to ensure preservation of ring connectivity and fused-ring integrity. If the scaffold fails either check, feature-agnostic MCS matching is used as a fallback, which ignores atomic identities (e.g., C vs. N) to preserve core topology across structurally similar compounds. R-groups (e.g., R_1_, R_2_, …, R_n_) are assigned to structural moieties not included in the core scaffold. To safeguard essential features, non-carbon atoms within the scaffold and mismatched atoms introduced by feature-agnostic MCS matching are replaced with generic placeholders such as Z or X.

### 2.2 Construction of an R-group Descriptor Library

We developed an extensive library of structure fragments, based on the analysis of 2,712 chemical patents. These patents were sourced from the European Patent Office (EPO) and United States Patent and Trademark Office (USPTO) through a CPC-based search using A61P, compound, and structure, which can quickly identify patents with specific therapeutic uses.

This library encompasses 246 curated R-group definitions, each corresponding to a standardized chemical concept. Each definition is linked to one or more representative SMILES strings that capture structural variations including tautomerism, stereochemistry, and substitution patterns observed in patents. In total, we extracted over 1,800 unique fragment types, defined as fundamental building blocks for Markush claim construction. Fragmentation was guided by bond context and functional group boundaries to avoid breaking conjugated systems, core rings, or reactive centers.

Fragments encompass functional groups, chain segments, and ring systems. Functional groups include electron-donating (e.g., hydroxyl, amino) and electron-withdrawing (e.g., cyano, sulfonyl) moieties. Chain fragments include straight and branched alkyls, described by IUPAC-based length ranges (e.g., C_1_ – C_3_, C_4_ – C_6_), while substitution patterns are expressed using relative terms such as “optionally substituted” or “1–3 substituents.” Ring systems encompass both aromatic and aliphatic mono- and bicyclic structures.

### 2.3 Expansion of R-group space via a fragment database

To enhance the scope of R-group representations and extend patent coverage, we have developed a comprehensive Fragment Library consisting of 10,490 distinct structural fragments. These fragments are sourced from various chemical databases, including BindingDB [25], DrugBank [26], PubChem [27], and ChEMBL [28]. The fragment structures are obtained using the molecular fragmentation tool, MacFrag [29], selected to represent common structural motifs observed in chemical patents.

For R-groups, those present in the R-Group Descriptor Library or composed of non-cyclic linear structures are selected and compared for structural similarity with fragments from the Fragment Library using precomputed Morgan fingerprints (ECFP), with a radius of 2, and the Tanimoto similarity coefficient. Although ECFP may be less accurate for small fragments, it provides a systematic and consistent basis for selecting R-group extensions based on structural similarity.

### 2.4 Hierarchical decomposition system of complex R-groups

For each labeled R-group, the corresponding structural fragments are obtained by automatically cleaving bonds at scaffold attachment points. For co-located R-groups sharing an identical attachment point, all fragments are grouped using cosine similarity on ECFP4 fingerprints to reduce downstream enumeration complexity.

After grouping, each R-group and its candidate fragments are processed sequentially. Fragments are matched against the R-Group Descriptor Library via SMILES comparison for validation. If a fragment is found in the library, its predefined description is directly output in standardized Markush notation (e.g., “C1–C4 alkyl”). For fragments not cataloged in the library, acyclic ones are resolved to generalized molecular formulas. Cyclic fragments undergo recursive processing, with structurally identical ring systems grouped and designated as a unified scaffold, and peripheral substituents are replaced with R-group descriptors.

## 3. Results

### 3.1 Evaluation

To evaluate the performance of SpaceExpander, we assembled an evaluation set of 24 publicly available chemical patent documents and extracted representative disclosed compounds from each patent as inputs. Four representative examples are provided in the Supplementary Information, and the input compound files for this evaluation are publicly available. For each patent, the extracted Markush scaffold was compared with the reference scaffold disclosed in the original patent by computing atomic-level accuracy, defined as the ratio of correctly recovered atoms to the total number of atoms in the reference scaffold. SpaceExpander achieved a mean atomic-level scaffold accuracy of 0.92, yielding identical scaffolds for 19 patents and only minor atomic differences in the remaining cases. For comparison, IntelliPatent was tested on the same dataset but could process only 2 of the 24 patents, with extracted scaffolds matching those from SpaceExpander.

### 3.2 Case Study: SpaceExpander Significantly Extends the Chemical Space Covered by the Osimertinib Patent

To assess the practical utility of SpaceExpander in real-world patent drafting, we conducted a case study using Osimertinib, a third-generation EGFR tyrosine kinase inhibitor (TKI). With strong clinical potential, Osimertinib rapidly became a key asset in the oncology landscape; however, multiple companies began developing alternative EGFR TKIs by modifying the molecular scaffold while maintaining biological activity [30-32].

We selected representative compounds from the Osimertinib patent (WO2013014448A1) and used them as input to draft a Markush claim. Figure 2 shows the chemical structures of all eight approved inhibitors, arranged by market approval date, with red highlights indicating structural differences. Notably, in the original Osimertinib patent, the structures of the seven sibling compounds were not included within the claim’s coverage. In contrast, the claim drafted by SpaceExpander successfully encompassed five compounds: Almonertinib, Mobocertinib, Befotertinib, Rezivertinib, and Rievirtonib. Lazertinib and Furmonertinib were not fully captured due to their distinct heterocyclic frameworks and side chain modifications.

**Fig. 2.**
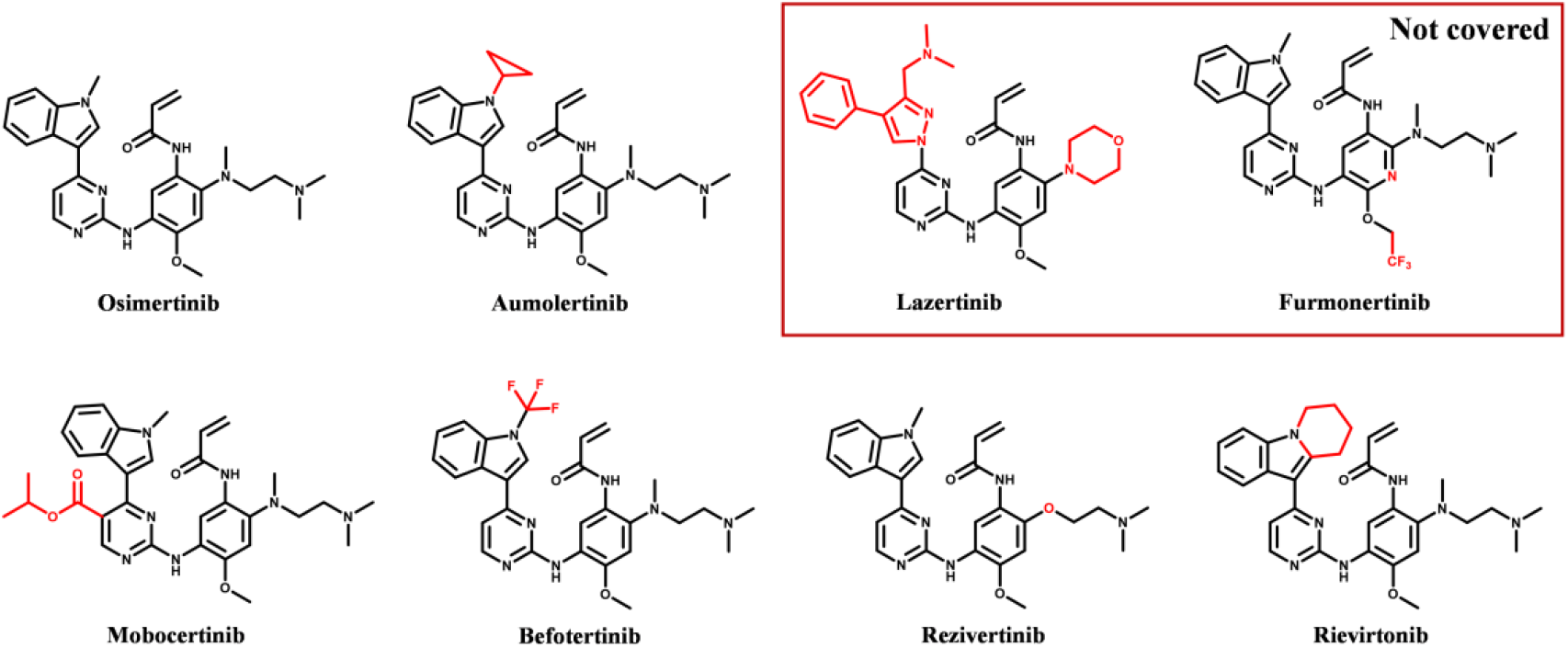
Structures of Osimertinib and other approved third-generation EGFR inhibitors, arranged by order of market approval. Differences between Osimertinib and the seven additional inhibitors are highlighted in red.

### 3.3 Case Study: SpaceExpander Outclasses IntelliPatent in Chemical Claim Generation

To evaluate the performance of SpaceExpander and IntelliPatent, we conducted a case study based on patent US6248771B1, which focuses on compounds designed to improve the oral bioavailability of hydrophobic drugs. Using the 17 example compounds disclosed in the patent, Figure 3 presents a comparison between the Markush formulas and R2 fragment descriptions.

**Fig. 3.**
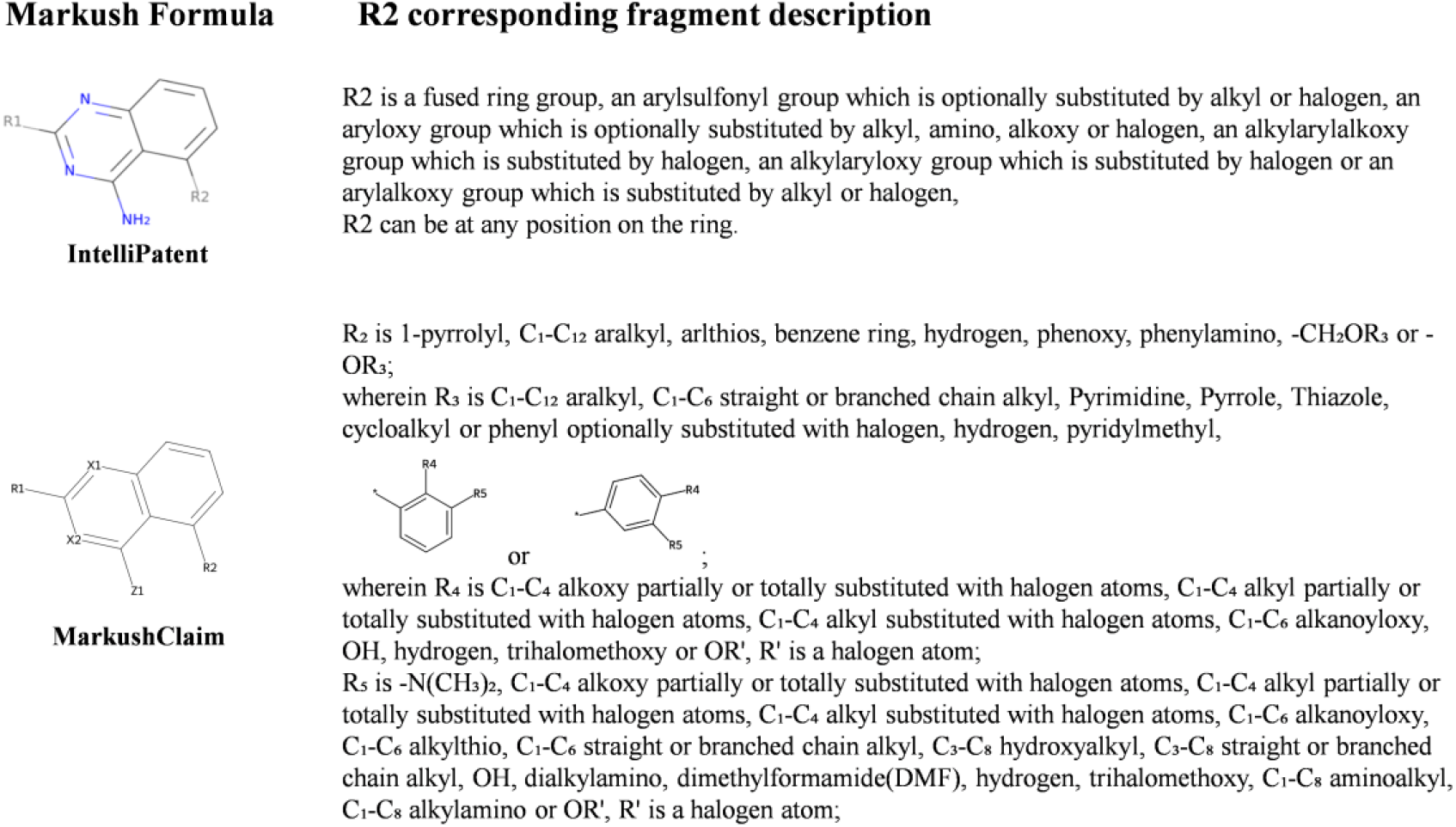
Comparison of the Markush structures and R_2_-group descriptions drafted by SpaceExpander and IntelliPatent for the compounds in patent ID US6248771B1.

In this case study, SpaceExpander provided a more detailed and chemically explicit handling of R-group descriptions than IntelliPatent. In particular, it preserved key structural features while refining R-group descriptions according to local structural context, rather than relying only on broader predefined descriptors.

### 3.4 Availability and deployment

We developed a user-friendly online server that allows users to upload compound files in either SDF format or CSV containing SMILES strings. Upon upload, the server processes the input and drafts a finalized claims document in Microsoft Word (.docx) format. Processing time depends on the number of compounds, typically ranging from a few minutes to over ten. Figure 4 shows the web interface, where users can submit files and initiate automated claim drafting.

**Fig. 4.**
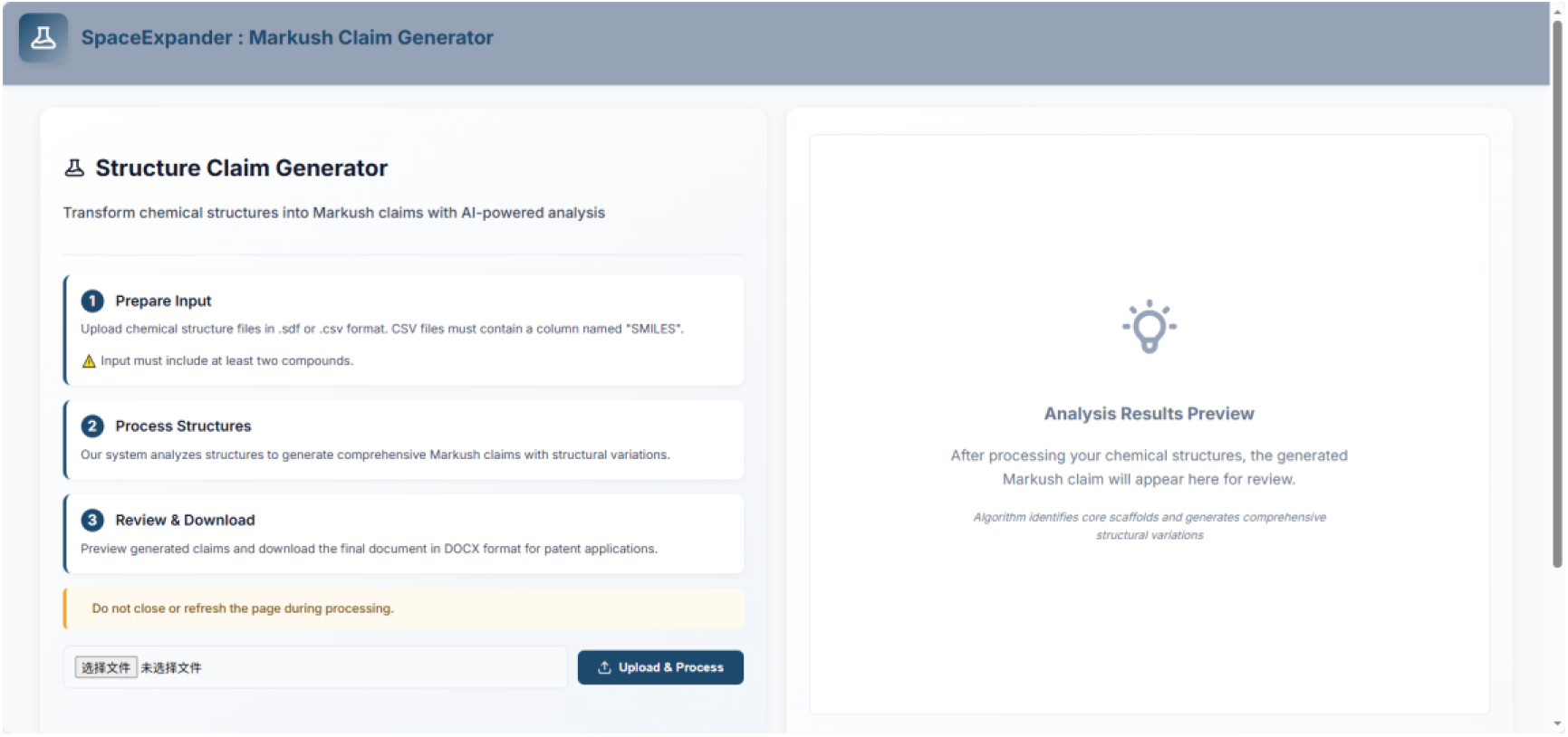
Web interface of the online implementation of SpaceExpander.

In response to concerns regarding data privacy and the risk of intellectual property leakage, especially when handling unpublished or confidential compound information, we also provide standalone executables for Windows, Linux, and macOS. These locally deployable versions retain the full functionality of the online platform while allowing users to process data entirely offline.

## 4. Conclusion

SpaceExpander provides a validated computational framework for automated Markush claim drafting and chemical space expansion. By integrating scaffold extraction, R-group decomposition, and substituent space expansion, the method enables the transformation of disclosed compounds into generalized draft claims while extending coverage of relevant chemical space.

Although the drafted outputs are supported by the evaluation results presented in this study, they may still differ from the refined language and legal precision of patents prepared by experienced professionals. Therefore, the drafted claims should be regarded as draft materials that require expert review and further legal revision before practical use. Nevertheless, the results demonstrate that SpaceExpander provides a useful method for automated Markush claim drafting and chemical space exploration. With continued advances in language generation and claim interpretation, frameworks of this kind may become increasingly valuable in supporting pharmaceutical patent drafting.

## Author contributions

Rui Wu developed the SpaceExpander method, implemented the algorithm, designed the web interface, and wrote the manuscript. Liyun Mao contributed to method development, data collection, and manuscript preparation. Yanyan Diao guided the research, evaluated the software, and assisted with manuscript revision. Honglin Li conceived the study and supervised the overall research. All authors reviewed and approved the final manuscript.

## Acknowledgments

This research received no external funding.

## Data Availability Statement

The source code of the SpaceExpander application is publicly available on GitHub at https://github.com/rwu527/SpaceExpander. A web-based version of SpaceExpander is freely accessible at https://www.lilab-ecust.cn/MarkushClaim, enabling online patent claim generation. For offline use, precompiled executables for Windows, Linux, and macOS are provided at https://drive.google.com/drive/folders/1DNeaFxnVlkqT0lW7YxIK2sOQMCxz6SkM?usp=drive_link.

