## Supplementary Information for "SpaceExpander: Automated Drafting and Evaluation of Markush Claims for Chemical Space Expansion"

We assembled an evaluation set of 24 publicly available chemical patent documents and extracted representative disclosed compounds from each patent as inputs to SpaceExpander. The extracted scaffolds were then compared with the reference scaffolds disclosed in the original patents using atomic-level accuracy. The input compound files for this evaluation are available at https://github.com/rwu527/SpaceExpander. For illustration, drafted editable claim texts for four representative patents from this 24-document evaluation set are presented below.

**Document 1: US 11 306 102 B2**

Claims

1. A compound of formula (I):

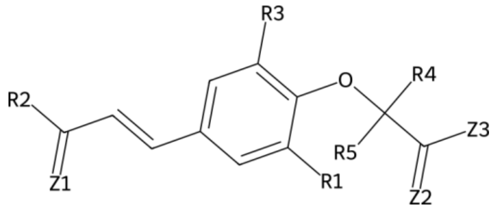

(I)

wherein:

$R_1$ is $-N(CH_3)_2$, $C_1$-$C_4$ alkoxy partially or totally substituted with halogen atoms, $C_1$-$C_4$ alkyl partially or totally substituted with halogen atoms, $C_1$-$C_6$ alkanoyl, $C_1$-$C_6$ alkanoyloxy, $C_1$-$C_6$ alkylthio, $C_1$-$C_6$ straight or branched chain alkyl, $C_3$-$C_8$ alkenyl, $C_3$-$C_8$ cycloalkyl, $C_3$-$C_8$ hydroxyalkyl, $C_3$-$C_8$ straight or branched chain alkyl, OH, a halogen atom, a straight or branched alkylene chain with 1 to 5 carbon atoms, dialkylamino, hydrogen, $C_1$-$C_8$ aminoalkyl, $C_1$-$C_8$ alkylamino or OR', R' is a halogen atom;

$R_3$ is $-N(CH_3)_2$, $C_1$-$C_4$ alkoxy partially or totally substituted with halogen atoms, $C_1$-$C_4$ alkyl partially or totally substituted with halogen atoms, $C_1$-$C_6$ alkanoyl, $C_1$-$C_6$ alkanoyloxy, $C_1$-$C_6$ alkylthio, $C_1$-$C_6$ straight or branched chain alkyl, $C_3$-$C_8$ alkenyl, $C_3$-$C_8$ cycloalkyl, $C_3$-$C_8$ hydroxyalkyl, $C_3$-$C_8$ straight or branched chain alkyl, OH, a halogen atom, a straight or branched alkylene chain with 1 to 5 carbon atoms, dialkylamino, hydrogen, $C_1$-$C_8$ aminoalkyl, $C_1$-$C_8$ alkylamino or OR', R' is a halogen atom;

$R_2$ is 4- to 7-membered ring, Benzene or

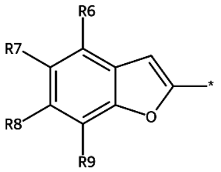

,

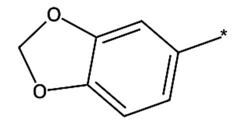

,

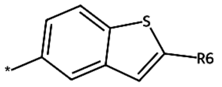

,

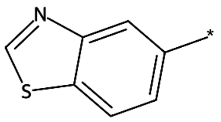

,

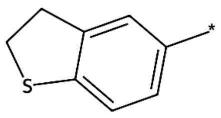

,

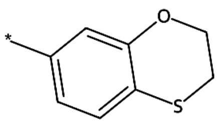

,

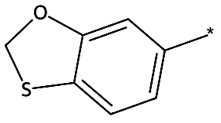

,

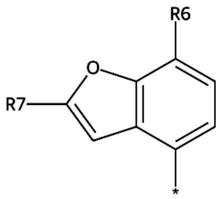

,

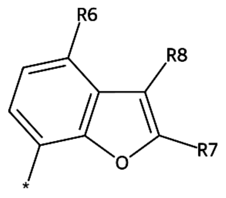

,

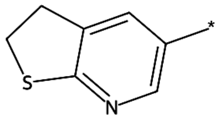
,
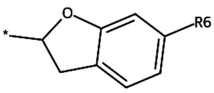
,
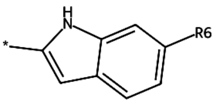
,
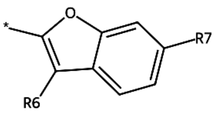
,
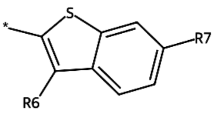
,
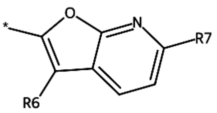
,
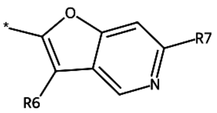
,
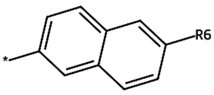
,
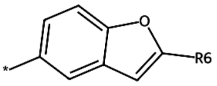
,
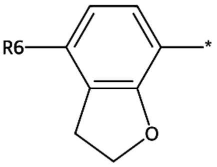
,
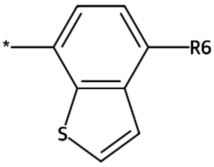
,
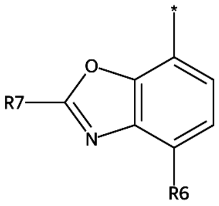
,
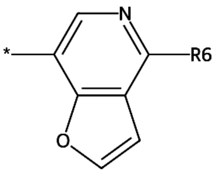
,
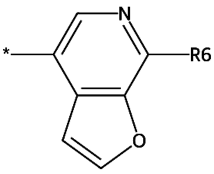
,
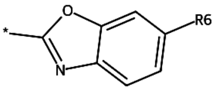
,
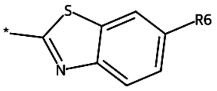
,
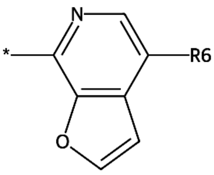
,
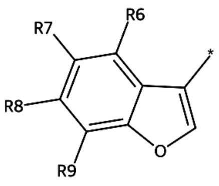
,
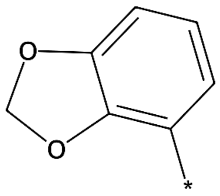
,
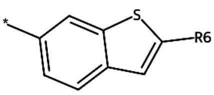
,
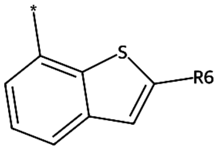
,

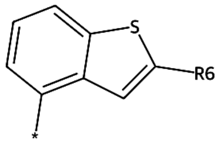
,
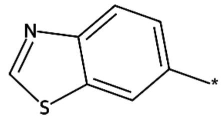
,
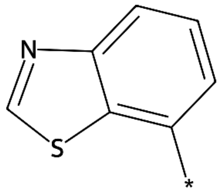
,
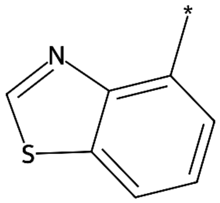
,
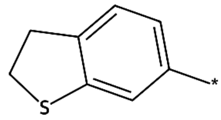
,
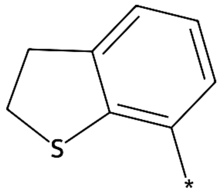
,
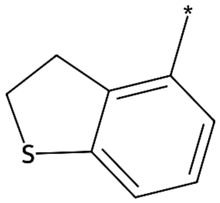
,
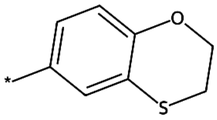
,
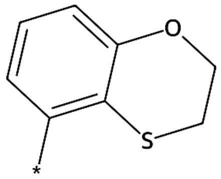
,
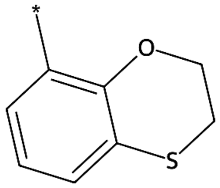
,
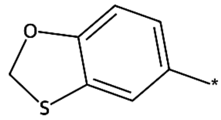
,
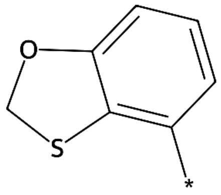
,
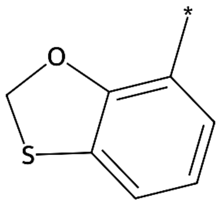
,
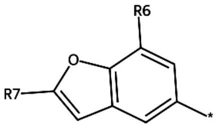
,
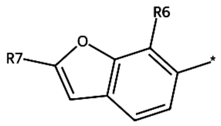
,
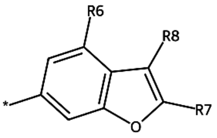
,
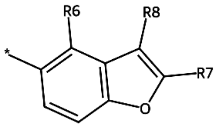
,
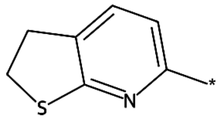
,
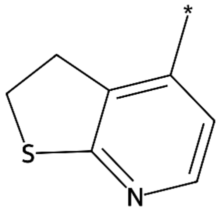
,
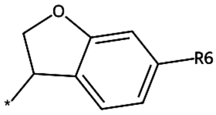
,
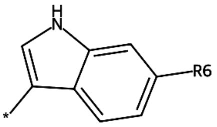
,

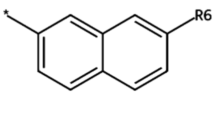

,

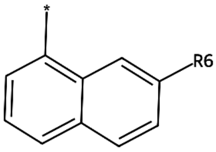

,

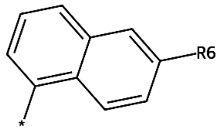

,

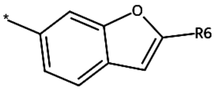

,

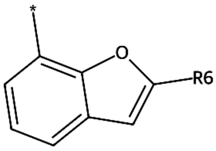

,

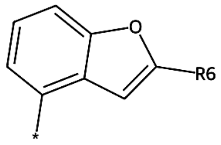

,

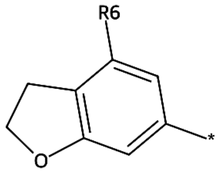

,

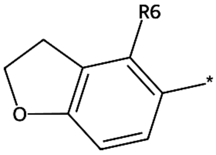

,

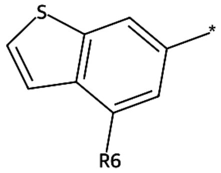

,

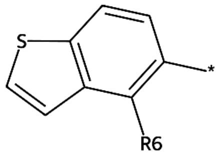

,

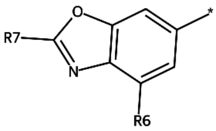

,

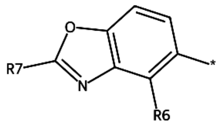

,

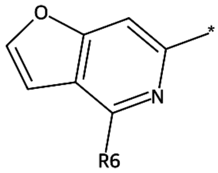

,

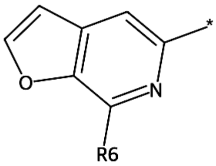

,

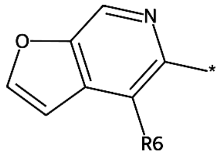

,

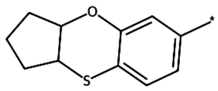

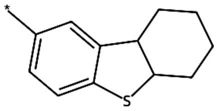

or ;

wherein $R_6$ is $-(CH_2)_2N(CH_3)_2$, $-(CH_2)_2OCH_3$, $-(CH_2)_3N(CH_3)_2$, $-(CH_2)_3OCH_3$, $-CH(CH_3)CH_2N(CH_3)_2$, $-CH(F)CH_2N(CH_3)_2$, $-CH(NH_2)CH_2N(CH_3)_2$, $-CH(OH)CH_2N(CH_3)_2$, $-CH_2C(=NH)OCH_3$, $-CH_2C(=S)N(CH_3)_2$, $-CH_2CH(CH_3)CH_2OCH_3$, $-CH_2CH(F)CH_2OCH_3$, $-CH_2CH(N(CH_3)_2)CH_3$, $-CH_2CH(NH_2)CH_2OCH_3$, $-CH_2CH(OH)CH_2OCH_3$, $-CH_2CH(OH)N(CH_3)_2$, $-CH_2COCH_2OCH_3$, $-CH_2N(C(=NH)CH_3)CH_3$, $-CH_2N(CHO)CH_3$, $-CH_2N(CH_3)CH(CH_3)_2$, $-CH_2N(CH_3)CH_2CH_3$, $-CH_2N(CH_3)_2$, $-CH_2N(COCH_3)CH_3$, $-CH_2NHCH_2OCH_3$, $-CH_2NHCH_3$, $-CH_2NHN(CH_3)_2$, $-CH_2OC(=NH)CH_3$, $-CH_2OCHO$, $-CH_2OCH_2CH_3$, $-CH_2OCH_2Cl$, $-CH_2OCH_2OCH_3$, $-CH_2OCH_3$, $-CH_2OCOOCH_3$, $-CH_2OF$, -

$CH_2OH$, $-CH_2ON(CH_3)_2$, $-CH_2ONHCH_3$, $-COCH_2N(CH_3)_2$, $-CON(CH_3)_2$, $-COOCH_3$, $-N(CH_3)CH_2OCH_3$, $-N(CH_3)_2$, $-NHCH_2N(CH_3)_2$, $-NHCH_2OCH_3$, $-OCH_2N(CH_3)_2$, $-OCH_3$, $-O_{13}CH_3$, $-SCH_2N(CH_3)_2$, $-SCH_2OCH_3$, $CH_3NHCO$, COOH, CHO or OH, $C_1$-$C_{12}$ straight-chain or branched acyl, $C_1$-$C_4$ alkoxy partially or totally substituted with halogen atoms, $C_1$-$C_4$ alkyl partially or totally substituted with halogen atoms, $C_1$-$C_6$ alkanoyl, $C_1$-$C_6$ alkanoyloxy, $C_1$-$C_6$ alkylthio, $C_1$-$C_6$ straight or branched chain alkyl, $C_1$-$C_8$ alkylsulfonyl, $C_3$-$C_8$ alkenyl, $C_3$-$C_8$ cycloalkyl, $C_3$-$C_8$ hydroxyalkyl, $C_3$-$C_8$ straight or branched chain alkyl, OH, a halogen atom, a straight or branched alkylene chain with 1 to 5 carbon atoms, alkylthioalkyl, carbamoyl, dialkylamino, dimethylamine, dimethylformamide(DMF), fluoralkoxy, hydrogen, substituted/unsubstituded aminos, thiols, trihalomethoxy, $C_1$-$C_8$ aminoalkyl, $C_1$-$C_8$ alkylamino or OR', R' is a halogen atom;

$R_7$ is $-N(CH_3)_2$, $C_1$-$C_4$ alkoxy partially or totally substituted with halogen atoms, $C_1$-$C_4$ alkyl partially or totally substituted with halogen atoms, $C_1$-$C_6$ alkanoyl, $C_1$-$C_6$ alkanoyloxy, $C_1$-$C_6$ alkylthio, $C_1$-$C_6$ straight or branched chain alkyl, $C_3$-$C_8$ cycloalkyl, $C_3$-$C_8$ hydroxyalkyl, $C_3$-$C_8$ straight or branched chain alkyl, OH, a halogen atom, a straight or branched alkylene chain with 1 to 5 carbon atoms, alkylthioalkyl, fluoralkoxy, hydrogen, thiols, trihalomethoxy, $C_1$-$C_8$ aminoalkyl, $C_1$-$C_8$ alkylamino or OR', R' is a halogen atom;

$R_8$ is $-CH_2OH$, $CH_3NHCO$, COO-alkyl, COOH, CHO or OH, $C_1$-$C_{12}$ straight-chain or branched acyl, $C_1$-$C_4$ alkoxy partially or totally substituted with halogen atoms, $C_1$-$C_4$ alkyl partially or totally substituted with halogen atoms, $C_1$-$C_4$ alkylamino, $C_1$-$C_4$ aminoalkyl, $C_1$-$C_6$ alkanoyl, $C_1$-$C_6$ alkanoyloxy, $C_1$-$C_6$ alkoxy, $C_1$-$C_6$ alkylthio, $C_3$-$C_8$ acyloxy, $C_3$-$C_8$ alkenoyl, $C_3$-$C_8$ alkoxycarbonyl, $C_3$-$C_8$ straight or branched chain alkyl, OH, a halogen atom, alkylthioalkyl, alkynyloxy, carbamoyl, fluoralkoxy, formyl, hydrogen, thiols or OR', R' is a halogen atom;

$R_9$ is $C_1$-$C_6$ alkylthio, hydrogen or thiols;

$R_4$ is $-N(CH_3)_2$, $C_1$-$C_4$ alkyl partially or totally substituted with halogen atoms, $C_1$-$C_4$ alkylamino, $C_1$-$C_6$ alkanoyl, $C_1$-$C_6$ straight or branched chain alkyl, $C_3$-$C_8$ cycloalkyl, a straight or branched alkylene chain with 1 to 5 carbon atoms or forms an unsubstituted/substituted $C_3$-$C_8$ cycloalkyl with $R_5$;

$R_5$ is $-N(CH_3)_2$, $C_1$-$C_4$ alkyl partially or totally substituted with halogen atoms, $C_1$-$C_4$ alkylamino, $C_1$-$C_6$ alkanoyl, $C_1$-$C_6$ straight or branched chain alkyl, $C_3$-$C_8$ cycloalkyl, a straight or branched alkylene chain with 1 to 5 carbon atoms or forms an unsubstituted/substituted $C_3$-$C_8$ cycloalkyl with $R_4$;

$Z_1$, $Z_2$ and $Z_3$ are the same or different and each independently represents C, N, S or O.

**Document 2：US 10 233 146 B2**

Claims

1. A compound of formula (I):

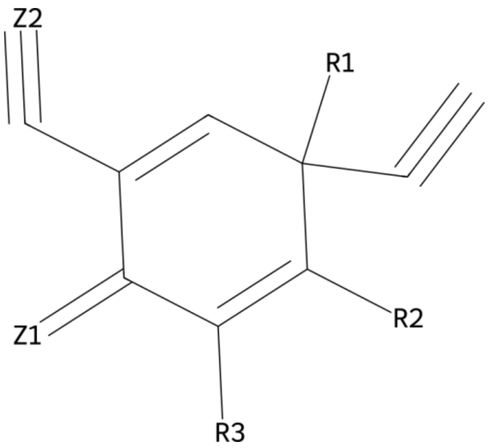

(I)

wherein:

$R_3$ is hydrogen or forms an unsubstituted/substituted benzene ring with $R_2$;

$R_1$ is -N($CH_3$)$_2$, 1,2-alkylvinylene, $C_1$-$C_4$ alkyl partially or totally substituted with halogen atoms, $C_1$-$C_4$ alkylamino, $C_1$-$C_6$ alkanoyl, $C_1$-$C_6$ straight or branched chain alkyl, $C_2$-$C_6$ alkenyl, $C_2$-$C_6$ alkynyl, $C_3$-$C_8$ alkenoyl, $C_3$-$C_8$ alkenyl, $C_3$-$C_8$ cycloalkyl, a straight, branched alkylene chain with 1 to 5 carbon atoms, alkynyloxy, hydrogen or forms the structure shown below:

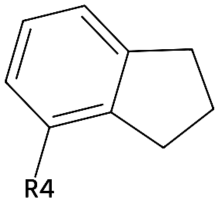

,

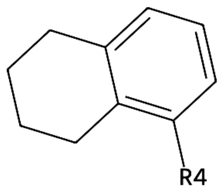

,

or a substituted $C_5$-$C_7$ cycloalkyl ring with $R_2$;

wherein $R_4$ is -N($CH_3$)$_2$, $C_1$-$C_4$ alkyl partially or totally substituted with halogen atoms, $C_1$-$C_4$ alkylamino, $C_1$-$C_6$ alkanoyl, $C_1$-$C_6$ straight or branched chain alkyl, $C_3$-$C_8$ cycloalkyl, a halogen atom, a straight or branched alkylene chain with 1 to 5 carbon atoms, hydrogen or OR', R' is a halogen atom;

$R_2$ is hydrogen, forms an unsubstituted/substituted benzene ring with $R_3$ or forms the structure shown below:

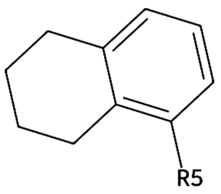

,

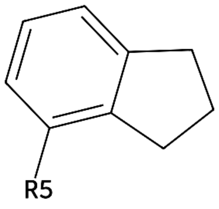

,

or a substituted $C_5$-$C_7$ cycloalkyl ring with $R_1$;

wherein $R_5$ is $-N(CH_3)_2$, $C_1$-$C_4$ alkyl partially or totally substituted with halogen atoms, $C_1$-$C_4$ alkylamino, $C_1$-$C_6$ alkanoyl, $C_1$-$C_6$ straight or branched chain alkyl, $C_3$-$C_8$ cycloalkyl, a halogen atom, a straight or branched alkylene chain with 1 to 5 carbon atoms, hydrogen or OR', R' is a halogen atom;

$Z_1$ represents C, N, S or O;

$Z_2$ represents C or N.

**Document 3: US 10 537 558 B2**

**Claims**

1. A compound of formula (I):

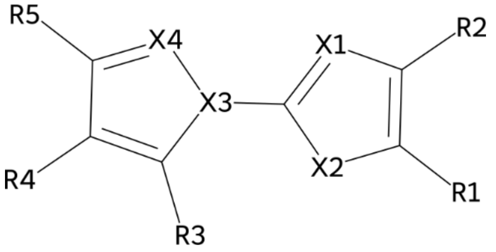

(I)

wherein:

$R_1$ is $-(CH_2)_2S(O)CH(CH_3)_2$, $-C(CH_3)(CH_3)CH(CH_3)_2$, $-C(F)(F)CH(CH_3)_2$, $-C(OH)(CH_3)CH(CH_3)_2$, $-C(S(O)CH_3)(CH_3)_2$, $-CH(CH_3)CH(NH_2)CH_3$, $-CH(CH_3)CH(SH)CH_3$, $-CH(CH_3)CH_2S(O)CH_3$, $-CH_2CH(CH_3)CH_2S(O)CH_3$, $-CH_2S(O)CH(CH_3)_2$, $-N(CH_3)(CH_3)CH(CH_3)_2$, $-N(CH_3)CH(CH_3)_2$, $-NCH(CH_3)_2$, $-NHS(O)CH(CH_3)_2$, $-ONHCH(CH_3)_2$, $-S(O)CH(CHO)CH_3$, $-S(O)CH(CH_3)CH_2CH_3$, $-S(O)CH(CH_3)_2$, $-S(O)CH_2CH(CH_3)_2$, $-SO_2CH(CH_3)_2$, $-SSCH(CH_3)_2$, $=C(CH_3)CH(CH_3)_2$, $=C(Cl)CH(CH_3)_2$, $=C(OH)CH(CH_3)_2$, $C_1$-$C_{12}$ straight-chain or branched acyl, $C_1$-$C_4$ alkyl partially or totally substituted with halogen atoms, $C_1$-$C_6$ alkanoyloxy, $C_1$-$C_6$ alkylthio, $C_1$-$C_6$ straight or branched chain alkyl, $C_3$-$C_8$ alkylamino, $C_3$-$C_8$ hydroxyalkyl, $C_3$-$C_8$ straight or branched chain alkyl, alkanesulphonyl, alkylthioalkyl, dialkylamino, hydrogen or $C_1$-$C_8$ aminoalkyl, $-SR_6$;

wherein $R_6$ is hydrogen, piperidine or $C_3$-$C_{10}$ cycloalkyl,

$R_2$ is $C_1$-$C_{12}$ aralkyl, $C_1$-$C_6$ straight or branched chain alkyl, $C_3$-$C_8$ cycloalkyl, Furan, Pyrimidine, Pyrrole, Thiazole, arlthios, aromatic, hetroaromatic which can be substituted with one, more halogeens, aromatic, hetroaromatic which can be substituted with one, more halogen atoms, benzene ring, cycloalkyl or phenyl optionally substituted with halogen, cycloalkyl or phenyl optionally substituted with hydroxyl or phenyl, cyclohexylene, furoyl, phenoxy, phenylamino, piperidine, pyrazine, pyridazine, pyridineamino, pyridylmethyl, $-C\equiv CR_7$ or

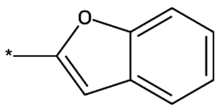

,

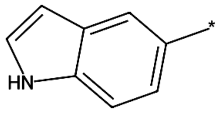

,

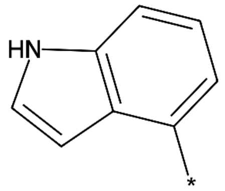

,

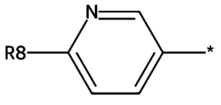

,

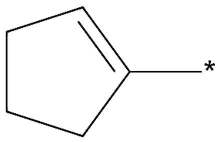

,

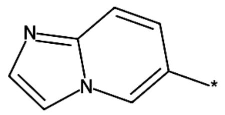

,

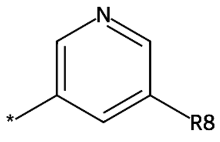
,
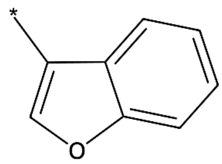
,
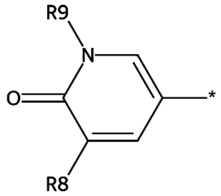
,
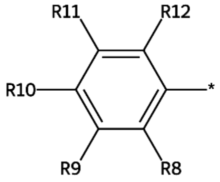
,
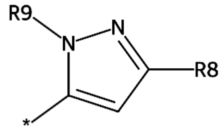
,
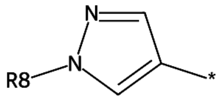
,
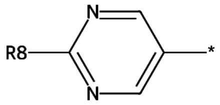
,
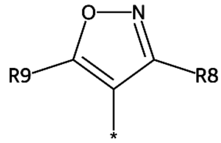
,
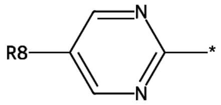
,
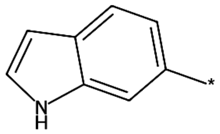
,
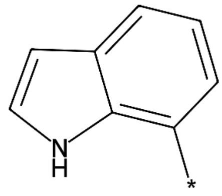
,
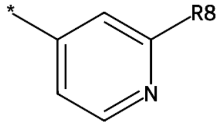
,
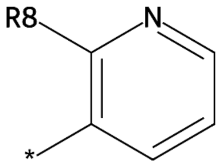
,
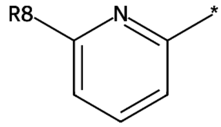
,
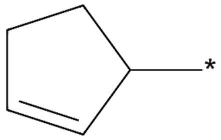
,
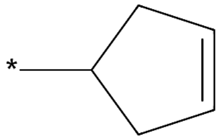
,
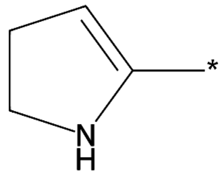
,
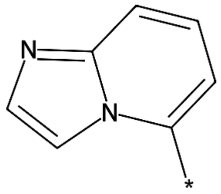
,
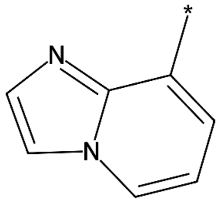
,
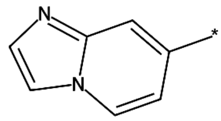
,
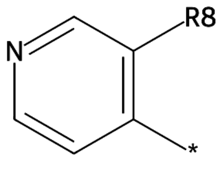
,

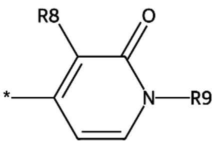

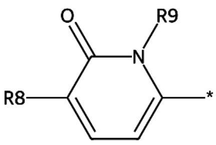

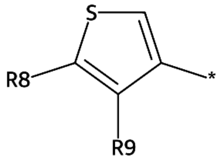

, , ,

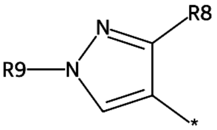

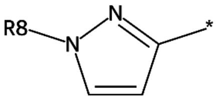

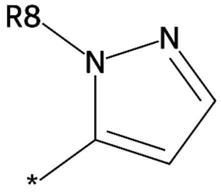

, , ,

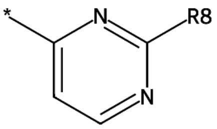

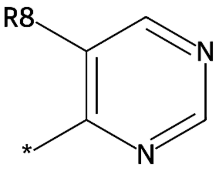

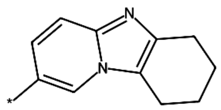

, , ,

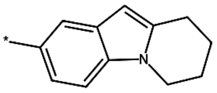

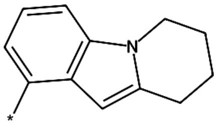

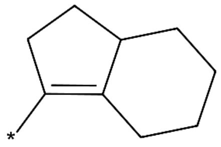

, or ;

wherein $R_7$ is $C_1$-$C_{12}$ aralkyl, $C_3$-$C_8$ cycloalkyl, Pyrimidine, arlthios, benzene ring, cycloalkyl or phenyl optionally substituted with hydroxyl or phenyl, hydrogen, phenoxy, phenylamino, piperidine or

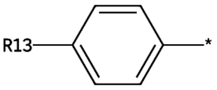

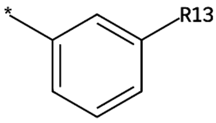

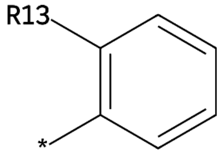

, , ,

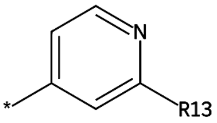

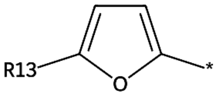

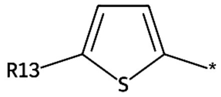

, , or

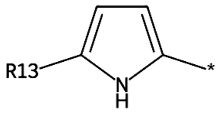

;

wherein $R_{13}$ is $C_1$-$C_4$ alkyl partially or totally substituted with halogen atoms, hydrogen or trihalomethoxy;

$R_8$ is $-N(CH_3)_2$, $C_1$-$C_4$ alkoxy partially or totally substituted with halogen atoms, $C_1$-$C_4$ alkyl partially or totally substituted with halogen atoms, $C_1$-$C_4$ alkylamino, $C_1$-$C_6$ alkanoyl, $C_1$-$C_6$ straight or branched chain alkyl, $C_1$-$C_8$ alkylsulfonyl, $C_3$-$C_8$ cycloalkyl, $NO_2$, Pyrimidine, a halogen atom, a straight or branched alkylene chain with 1 to 5 carbon atoms, benzene ring, cycloalkyl or phenyl optionally substituted with halogen, dimethylamine, fluoralkoxy, hydrogen, trihalomethoxy or OR', R' is a halogen atom;

$R_9$ is $-N(CH_3)_2$, $C_1$-$C_4$ alkoxy partially or totally substituted with halogen atoms, $C_1$-$C_4$ alkyl partially or totally substituted with halogen atoms, $C_1$-$C_4$ alkylamino, $C_1$-$C_6$ alkanoyl, $C_1$-$C_6$ straight or branched chain alkyl, $C_3$-$C_8$ cycloalkyl, a halogen atom, a straight or branched alkylene chain with 1 to 5 carbon atoms, fluoralkoxy, hydrogen, trihalomethoxy or OR', R' is a halogen atom;

$R_{10}$ is $-(CH_2)_2C(F)(F)CH_2CH_3$, $-(CH_2)_2C(F)(F)CH_2OH$, $-(CH_2)_2CONH(CH_2)_2OCH_3$, $-(CH_2)_2OCH_3$, $-(CH_2)_2S(F)(F)(F)(F)F$, $-C(F)(CH_2CH_3)CH_2CH_3$, $-C(F)(F)(CH_2)_2CH_3$, $-C(F)(F)CH(CH_3)_2$, $-C(F)(F)CH(F)F$, $-C(F)(F)CH_2CH=CH_2$, $-C(F)(F)CH_2CHO$, $-C(F)(F)CH_2CH_3$, $-C(F)(F)CH_2F$, $-C(F)(F)CH_2NH_2$, $-C(F)(F)CH_2OCH_3$, $-C(F)(F)CH_2OH$, $-C(F)(F)CH_3$, $-C(F)(F)COCH_3$, $-C(F)(F)Cl$, $-C(F)(F)OH$, $-CF_3$, $-CH(CH_3)CF_3$, $-CH(F)CF_3$, $-CH(F)F$, $-CH_2C(F)(F)CH_2CH_3$, $-CH_2CF_3$, $-CH_2CH(F)CH_3$, $-CH_2COCH_2OCH_3$, $-CH_2CONH(CH_2)_2OCH_3$, $-CH_2NHCOOCH_3$, $-CH_2O(CH_2)_2COOH$, $-CH_2S(F)(F)(F)(F)F$, $-CO(CH_2)_2OCH_3$, $-CONH(CH_2)_2Br$, $-CONH(CH_2)_2CH_3$, $-CONH(CH_2)_2Cl$, $-CONH(CH_2)_2F$, $-CONH(CH_2)_2NH_2$, $-CONH(CH_2)_2OCHO$, $-CONH(CH_2)_2OCH_2CH_3$, $-CONH(CH_2)_2OCH_3$, $-CONH(CH_2)_2SH$, $-CONH(CH_2)_3OCH_3$, $-CONHCH_2Br$, $-CONHCH_2CH=NH$, $-CONHCH_2CHO$, $-CONHCH_2COOCH_3$, $-CONHCH_2OCH_3$, $-CONHCH_2SCH_3$, $-CONHCOCH_2OCH_3$, $-COO(CH_2)_2CH_3$, $-COOCH_2OCH_3$, $-N(CH_3)_2$, $-NHCF_3$, $-NHCH_2NHCHO$, $-NHSO_2CH_3$, $-OBe_2(F)(F)F$, $-OC(F)(F)CH_2CH_3$, $-OCF_3$, $-OCH(F)F$, $-OCH_2CH(F)F$, $-OCH_2NHCHO$, $-S(F)(F)(F)(F)F$, $-SCF_3$, $=C(F)CF_3$, $=C(I)CF_3$, $=C(OH)CF_3$, $=CHCF_3$, $C_1$-$C_4$ alkoxy partially or totally substituted with halogen atoms, $C_1$-$C_4$ alkyl partially or totally substituted with halogen atoms, $C_1$-$C_4$ alkylamino, $C_1$-$C_6$ alkanoyl, $C_1$-$C_6$ alkanoyloxy, $C_1$-$C_6$ straight or branched chain alkyl, $C_3$-$C_8$ cycloalkyl, OH, a halogen atom, a straight or branched alkylene chain with 1 to 5 carbon atoms, alkanesulphonyl, dimethylformamide(DMF), hydrogen, methylethylcarbamoyl, trihalomethoxy or OR', R' is a halogen atom;

$R_{11}$ is $-(CH_2)_2CN$, $-BHCN$, $-CH(Br)CN$, $-CH(CN)OH$, $-CH(COOCH_3)CH_3$, $-CH(OCOCH_3)CH_3$, $-CH_2CH(OC(=S)S)CH_3$, $-CH_2CH(OC(=S)SH)CH_3$, $-CH_2CN$, $-CH_2COCH(CH_3)_2$, $-CH_2COOCH(CH_3)_2$, $-CN$, $-COCH(Br)CH_3$, $-COCH(OCH_3)CH_3$, $-COCH(SH)CH_3$, $-COCH_2CH(CH_3)_2$, $-COCN$, $-CONHCH(CH_3)_2$, $-COOCH(CH_3)_2$, $-COOCH(Cl)CH_3$, $-COOCH(F)CH_3$, $-COOCH(F)F$, $-COOCH_2CH_3$, $-COOCH_3$, $-COSCH(CH_3)_2$, $-N(CH_3)_2$, -

$N(CN)CH_3$, -NCN, -NHCN, -OCN, $-OCOCH(CH_3)_2$, -SCN, $=C(CN)NH_2$, =C(CN)OH, =CHCN, CONH-alkyl, $C_1$-$C_4$ alkoxy partially or totally substituted with halogen atoms, $C_1$-$C_4$ alkyl partially or totally substituted with halogen atoms, $C_1$-$C_4$ alkylamino, $C_1$-$C_6$ alkanoyl, $C_1$-$C_6$ alkanoyloxy, $C_1$-$C_6$ straight or branched chain alkyl, $C_3$-$C_8$ cycloalkyl, OH, a halogen atom, a straight or branched alkylene chain with 1 to 5 carbon atoms, fluoralkoxy, hydrogen or OR', R' is a halogen atom;

$R_{12}$ is $-(CH_2)_2CN$, -BHCN, -CH(Br)CN, -CH(CN)OH, $-CH(COOCH_3)CH_3$, $-CH(OCOCH_3)CH_3$, $-CH_2CH(OC(=S)S)CH_3$, $-CH_2CH(OC(=S)SH)CH_3$, $-CH_2CN$, $-CH_2COCH(CH_3)_2$, $-CH_2COOCH(CH_3)_2$, -CN, $-COCH(Br)CH_3$, $-COCH(OCH_3)CH_3$, $-COCH(SH)CH_3$, $-COCH_2CH(CH_3)_2$, -COCN, $-CONHCH(CH_3)_2$, $-COOCH(CH_3)_2$, $-COOCH(Cl)CH_3$, $-COOCH(F)CH_3$, -COOCH(F)F, $-COOCH_2CH_3$, $-COOCH_3$, $-COSCH(CH_3)_2$, $-N(CH_3)_2$, $-N(CN)CH_3$, -NCN, -NHCN, -OCN, $-OCOCH(CH_3)_2$, -SCN, $=C(CN)NH_2$, =C(CN)OH, =CHCN, $C_1$-$C_4$ alkoxy partially or totally substituted with halogen atoms, $C_1$-$C_4$ alkyl partially or totally substituted with halogen atoms, $C_1$-$C_4$ alkylamino, $C_1$-$C_4$ aminoalkyl, $C_1$-$C_6$ alkanoyl, $C_1$-$C_6$ alkanoyloxy, $C_1$-$C_6$ straight or branched chain alkyl, $C_1$-$C_8$ alkylsulfonyl, $C_3$-$C_8$ cycloalkyl, OH, a halogen atom, a straight or branched alkylene chain with 1 to 5 carbon atoms, carbamoyl, dimethylamine, fluoralkoxy, hydrogen, substituted/unsubstituded aminos or OR', R' is a halogen atom;

$R_3$ is $-(CH_2)_2CON(CH_3)(CH_2)_2OH$, $-(CH_2)_2CON(CH_3)(CH_2)_3NH_2$, $-(CH_2)_2CONH(CH_2)_2OCH_3$, $-(CH_2)_2CONHOCH_3$, $-(CH_2)_2CONHSO_2(CH_2)_2CH_3$, $-(CH_2)_2OCHO$, $-(CH_2)_2OCH_3$, $-(CH_2)_2SO_2CH_3$, $-(CH_2)_2SO_2F$, $-(CH_2)_2SO_2O$, $-(CH_2)_2SO_2OH$, $-(CH_2)_3N(NH_2)CH_3$, $-(CH_2)_3SO_2OH$, $-CH(CH_3)(CH_2)_3NH_2$, $-CH_2COCH_2OCH_3$, $-CH_2CON(CH_3)(CH_2)_2OH$, $-CH_2CON(CH_3)(CH_2)_3NH_2$, $-CH_2CONH(CH_2)_2OCH_3$, $-CH_2CONHOCH_3$, $-CH_2CONHSO_2(CH_2)_2CH_3$, $-CH_2N(CHO)CH_3$, $-CH_2N(CH_3)(CH_2)_2NH_2$, $-CH_2N(CH_3)(CH_2)_2OH$, $-CH_2N(CH_3)(CH_2)_3NH_2$, $-CH_2N(CONH_2)CH_3$, $-CH_2N(OH)CHO$, $-CH_2NH(CH_2)_3NH_2$, $-CH_2NHCOOCH_3$, $-CH_2O(CH_2)_2COOH$, $-CH_2OCH_2CHO$, $-CH_2OH$, $-CH_2ONHCHO$, $-CH_2SO_2CH_2CH_3$, $-CH_2SO_2NHCH_3$, $-CO(CH_2)_2OCH_3$, $-CO(CH_2)_3CH_3$, $-CO(CH_2)_3NH_2$, $-CO(CH_2)_3OH$, $-COCH_2NHCH_2CH_3$, $-COCH_2NHOCH_3$, $-COCH_3$, $-CON(CHO)(CH_2)_2OH$, $-CON(CHO)(CH_2)_3NH_2$, $-CON(CHO)CH_3$, $-CON(CH_2CH_3)(CH_2)_2OH$, $-CON(CH_2CH_3)(CH_2)_3NH_2$, $-CON(CH_3)(CH_2)_2CONH_2$, $-CON(CH_3)(CH_2)_2OH$, $-CON(CH_3)(CH_2)_3NH_2$, $-CON(CH_3)(CH_2)_3OH$, $-CON(CH_3)(CH_2)_4NH_2$, $-CON(CH_3)CH_2CH_3$, $-CON(CH_3)CH_2COCH_2NH_2$, $-CON(CH_3)CH_2COOH$, $-CON(CH_3)CH_2OH$, $-CON(CH_3)_2$, $-CON(CO(CH_2)_2NH_2)CH_3$, $-CON(COCH_2OH)CH_3$, $-CON(NH_2)CH_3$, $-CON(OH)CH_2CH_3$, $-CON(OH)CH_3$, $-CONH(CH_2)_2Br$, $-CONH(CH_2)_2CH_3$, $-CONH(CH_2)_2Cl$, $-CONH(CH_2)_2F$, $-CONH(CH_2)_2NH_2$, $-CONH(CH_2)_2O(CH_2)_2O(CH_2)_2O(CH_2)_2N_3$, $-CONH(CH_2)_2OCHO$, $-CONH(CH_2)_2OCH_2CH_3$, $-CONH(CH_2)_2OCH_3$, $-CONH(CH_2)_2SH$, $-CONH(CH_2)_3OCH_3$, $-CONHCH_2Br$, $-CONHCH_2CH=CH_2$, $-CONHCH_2CH=NH$, $-CONHCH_2CHO$, $-CONHCH_2CH_3$, $-CONHCH_2CN$, $-CONHCH_2COOCH_3$, $-CONHCH_2C\equiv CH$, $-CONHCH_2OCH_3$, $-CONHCH_2OH$, $-CONHCH_2SCH_3$, $-CONHCOCH_2OCH_3$, $-CONHNHCH_2CH_3$, -CONHO, -CONHOCHO, $-CONHOCH_2CH_3$, $-CONHOCH_3$, $-CONHSO_2(CH_2)_2CHO$, $-CONHSO_2(CH_2)_2CH_3$, $-CONHSO_2(CH_2)_3CH_3$, $-CONHSO_2CH_2COCH_3$, $-CONHSO_2COCH_2CH_3$, $-COO(CH_2)_2CH_3$, $-COO(CH_2)_2NH_2$, $-COO(CH_2)_2OH$, $-COOCH_2OCH_3$, $-COOCH_3$, $-COONHCH_3$, $-COOOCH_3$, $-COS(CH_2)_2CH_3$, $-COS(CH_2)_2OH$, $-COSO_2CH_3$, $-COSO_2H$, $-N(CHO)(CH_2)_2OH$, $-N(CH_2CH_3)(CH_2)_2NH_2$, $-N(CH_2CH_3)(CH_2)_2OH$, $-N(CH_3)(CH_2)_2NH_2$, $-N(CH_3)(CH_2)_3NH_2$, -

$N(CH_3)(CH_2)_3OH$, $-N(COCH_2NH_2)CH_3$, $-N(COCH_2OH)CH_3$, $-NHC(=NH)NHOCH_3$, $-NHC(=S)NHOCH_3$, $-NHCH_2NHCHO$, $-NHCO(CH_2)_2CH_3$, $-NHCONHOCH_3$, $-NHN(CHO)CH_3$, $-NHSO_2CH_2CH_3$, $-NHSO_2CH_2F$, $-O(CH_2)_2CHO$, $-OCH_2NHCHO$, $-OCOCH_3$, $-OCONHOCH_3$, $-ON(CHO)CH_3$, $-ONHCOCH_3$, $-ONHCOOCH_3$, $-SCH_2NHCHO$, $-SO_2(CH_2)_2CH_3$, $=CHCONHOCH_3$, COO-alkyl, COOH, CHO or OH, $C_1$-$C_4$ alkyl partially or totally substituted with halogen atoms, $C_1$-$C_6$ alkanoyl, carbamoyl, dimethylformamide(DMF), formyl, hydrogen or methylethylcarbamoyl, $-COR_{14}$;

wherein $R_{14}$ is a straight, branched alkylene chain with 1 to 5 carbon atoms or

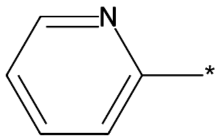
,
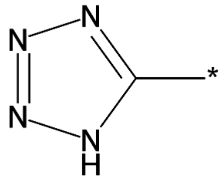
,
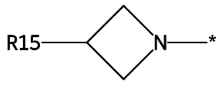

,
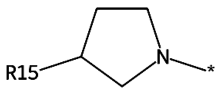

,
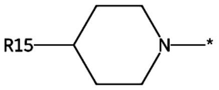

,
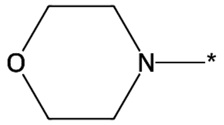

,
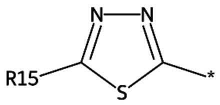

,
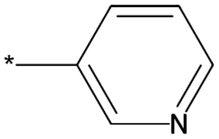

,
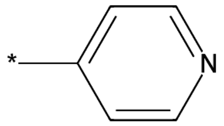

,
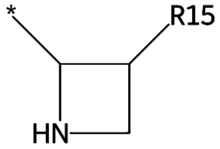

,
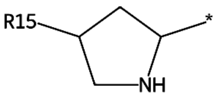

,
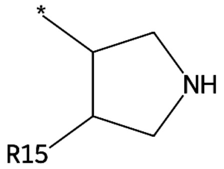

,
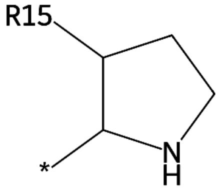

,
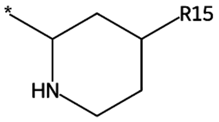

,
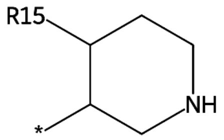

,

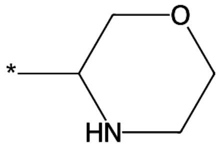 , 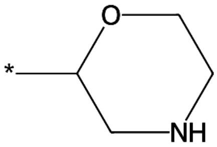 , 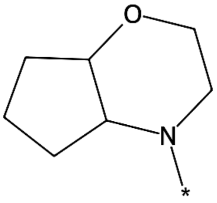 or

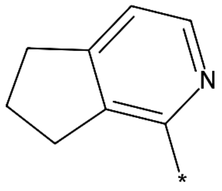 ;

wherein $R_{15}$ is $-N(C_2H_5)_2$, $C_1$-$C_4$ alkyl partially or totally substituted with halogen atoms, amino $C_1$-$C_4$ alkyl, carbamoyl, dialkylamino, hydrogen, substituted/unsubstituded aminos, trihalomethoxy or $C_1$-$C_8$ aminoalkyl;

$R_4$ is $C_1$-$C_{12}$ aralkyl, $C_1$-$C_4$ alkyl partially or totally substituted with halogen atoms, $C_1$-$C_6$ straight or branched chain alkyl, $C_3$-$C_8$ cycloalkyl, Indole, Isoquinoline, Pyrimidine, Pyrrole, a halogen atom, arlthios, aromatic, hetroaromatic which can be substituted with one, more halogen atoms, benzene ring, cycloalkyl or phenyl optionally substituted with halogen, cycloalkyl or phenyl optionally substituted with hydroxyl or phenyl, hydrogen, phenoxy, phenylamino, piperidine, pyrazine, pyridazine, pyridineamino, pyridylmethyl, $-CH_2R_{16}$ or

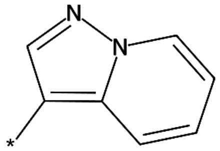 , 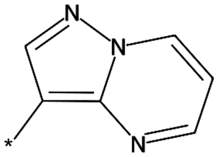 , 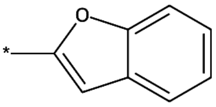 ,

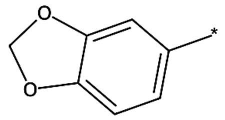 , 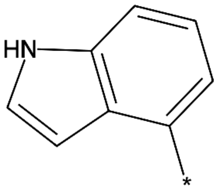 , 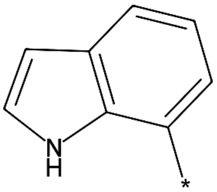 ,

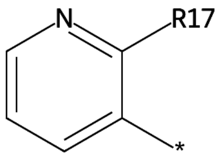 , 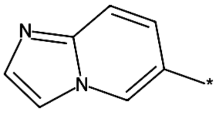 , 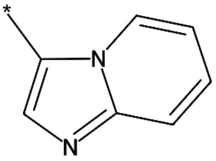 ,

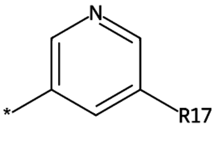
,
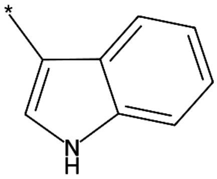
,
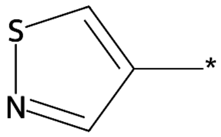
,
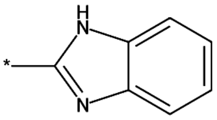
,
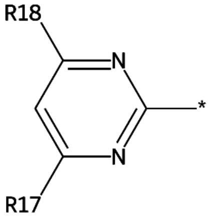
,
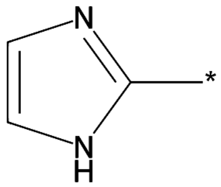
,
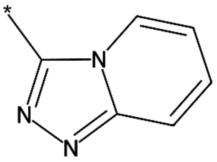
,
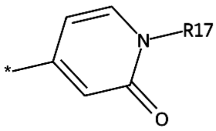
,
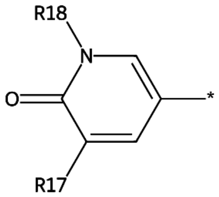
,
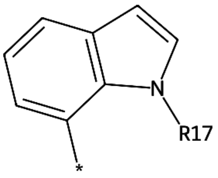
,
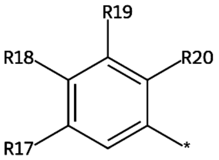
,
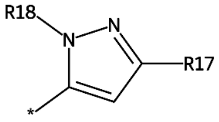
,
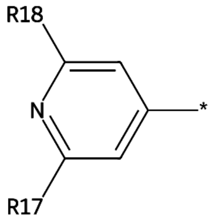
,
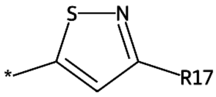
,
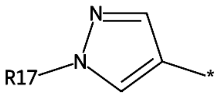
,
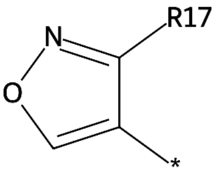
,
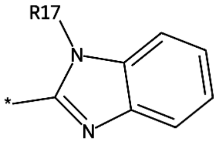
,
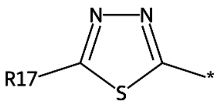
,
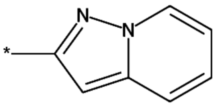
,
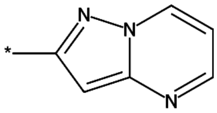
,
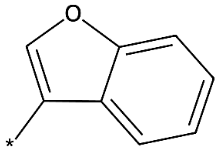
,

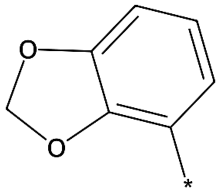
,
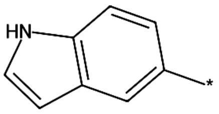
,
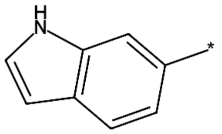
,
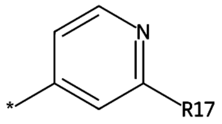
,
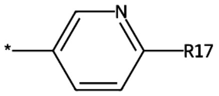
,
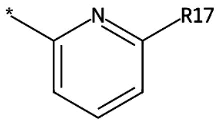
,
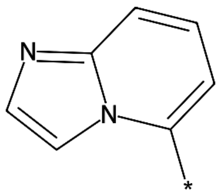
,
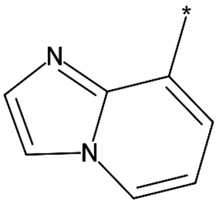
,
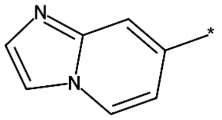
,
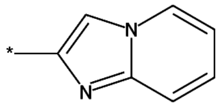
,
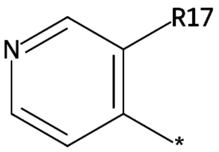
,
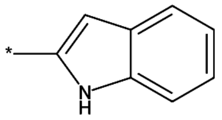
,
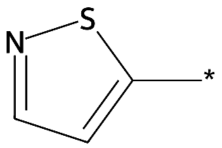
,
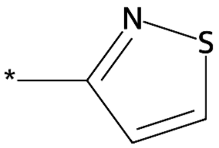
,
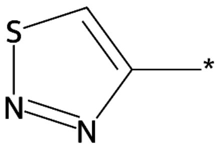
,
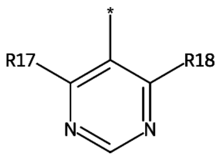
,
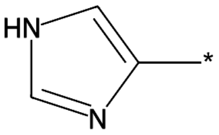
,
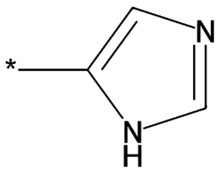
,
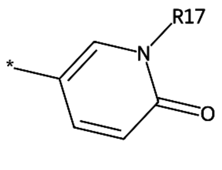
,
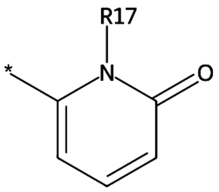
,
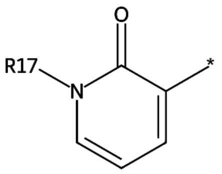
,

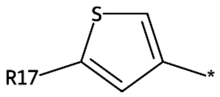
,
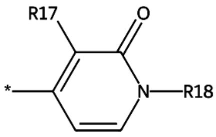
,
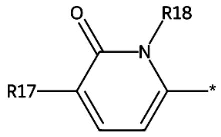
,
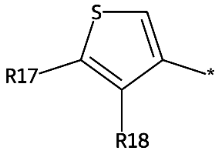
,
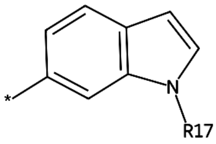
,
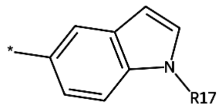
,
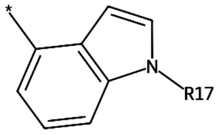
,
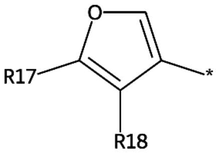
,
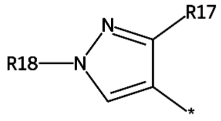
,
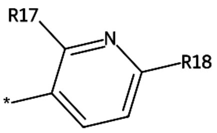
,
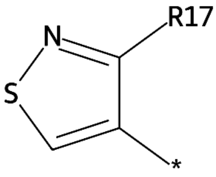
,
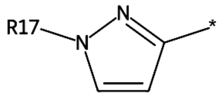
,
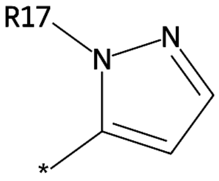
,
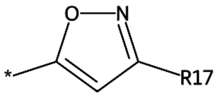
,
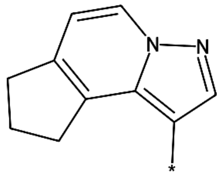
,
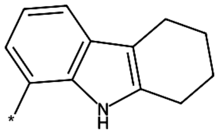
,
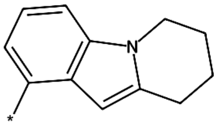
,
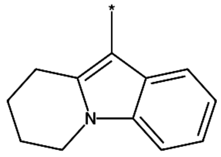
or
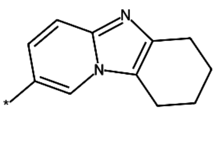
;

wherein $R_{16}$ is $C_1$-$C_{12}$ aralkyl, Pyrimidine, arlthios, benzene ring, hydrogen, phenoxy, phenyl substituted by 1-2 of amino, phenylamino, pyridineamino or

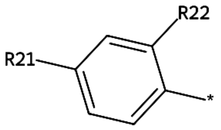

,

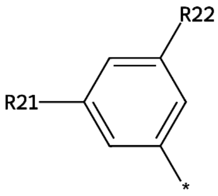

or

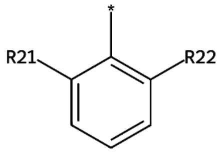

;

wherein $R_{21}$ is $-CH_2OH$, COOH, CHO or OH, $C_1$-$C_4$ alkyl partially or totally substituted with halogen atoms, hydrogen or trihalomethoxy;

$R_{22}$ is $-(CH_2)_2NHSO_2N(CH_3)_2$, $-CH(CH_3)SO_2CH_3$, $-CH_2N(CH_3)SO_2OH$, $-CH_2NHSO_2N(CH_3)_2$, $-CH_2OH$, $-CH_2SO_2NH_2$, $-N(CH_3)SO_2CH_3$, $-N(CH_3)SO_2OH$, $-NHCH_2SO_2CH_3$, $-NHCON(CH_3)_2$, $-NHSO(=CH_2)CH_3$, $-NHSO_2CH=CH_2$, $-NHSO_2CH_2Br$, $-NHSO_2CH_2CH_3$, $-NHSO_2CH_2Cl$, $-NHSO_2CH_2F$, $-NHSO_2CH_3$, $-NHSO_2N(CH_3)_2$, $-NHSO_2NHCH_3$, $-NHSO_2NH_2$, $-NHSO_2OCH_3$, $-NHSO_2OH$, $-SO_2CH(CH_3)_2$, $-SO_2CH_3$, COOH, CHO or OH, $C_1$-$C_4$ alkyl partially or totally substituted with halogen atoms, $C_3$-$C_8$ aminoalkyl, $NO_2$, alkanesulphonyl, amino $C_1$-$C_4$ alkyl, hydrogen or trihalomethoxy;

$R_{17}$ is $-(CH_2)_2CN$, $-(CH_2)_2NHOCH_3$, $-(CH_2)_2OCH(CH_3)_2$, $-(CH_2)_2OCHO$, $-(CH_2)_2OCH_2CH_3$, $-(CH_2)_2OCH_2NH_2$, $-(CH_2)_2OCH_2OH$, $-(CH_2)_2OCH_3$, $-(CH_2)_2OCN$, $-(CH_2)_2ON(CH_3)_2$, $-(CH_2)_2ONH_2$, $-(CH_2)_2OOH$, $-(CH_2)_2SOCH_3$, $-(CH_2)_3OCH_3$, $-(CH_2)_4OCH_3$, -BHCN, -CH(Br)CN, $-CH(CH_3)(CH_2)_2OCH_3$, -CH(CN)OH, $-CH(COOCH_3)CH_3$, $-CH(NH_2)(CH_2)_2OCH_3$, $-CH(OCOCH_3)CH_3$, $-CH_2CH(OC(=S)S)CH_3$, $-CH_2CH(OC(=S)SH)CH_3$, $-CH_2CN$, $-CH_2COCH(CH_3)_2$, $-CH_2COOCH(CH_3)_2$, $-CH_2COOCH_3$, $-CH_2O(CH_2)_2OCH_3$, $-CH_2O(CH_2)_3NH_2$, $-CH_2OCH_3$, $-CH_2OH$, -CN, $-CO(CH_2)_2OCH_3$, $-COCH(Br)CH_3$, $-COCH(OCH_3)CH_3$, $-COCH(SH)CH_3$, $-COCH_2CH(CH_3)_2$, $-COCH_2OCH_3$, -COCN, $-CONHCH(CH_3)_2$, $-COOCH(CH_3)_2$, $-COOCH(Cl)CH_3$, $-COOCH(F)CH_3$, -COOCH(F)F, $-COOCH_2CH_3$, $-COOCH_3$, $-COSCH(CH_3)_2$, $-N(CH_3)(CH_2)_2OCH_3$, $-N(CH_3)_2$, $-N(CN)CH_3$, -NCN, $-NH(CH_2)_2OCH_3$, -NHCN, -OCN, $-OCOCH(CH_3)_2$, -SCN, $=C(CN)NH_2$, =C(CN)OH, $=CH(CH_2)_2OCH_3$, =CHCN, COOH, CHO or OH, $C_1$-$C_{12}$ straight-chain or branched acyl, $C_1$-$C_4$ alkoxy partially or totally substituted with halogen atoms, $C_1$-$C_4$ alkyl partially or totally substituted with halogen atoms, $C_1$-$C_4$ alkylamino, $C_1$-$C_6$ alkanoyl, $C_1$-$C_6$ alkanoyloxy, $C_1$-$C_6$ alkoxy, $C_1$-$C_6$ straight or branched chain alkyl, $C_1$-$C_8$ alkylsulfonyl, $C_3$-$C_8$ alkoxycarbonyl, $C_3$-$C_8$ aminoalkyl, $C_3$-$C_8$ cycloalkyl, $C_3$-$C_8$ hydroxyalkyl, $C_3$-$C_8$ straight or branched chain alkyl, OH, a halogen atom, a straight or branched alkylene chain with 1 to 5 carbon atoms, alkylthioalkyl, dimethylamine, dimethylformamide(DMF), fluoralkoxy, hydrogen, methylethylcarbamoyl, trihalomethoxy or OR', R' is a halogen atom;

$R_{18}$ is $-N(CH_3)_2$, $CH_3NHCO$, $C_1$-$C_4$ alkoxy partially or totally substituted with halogen atoms, $C_1$-$C_4$ alkyl partially or totally substituted with halogen atoms, $C_1$-$C_4$ alkylamino, $C_1$-$C_6$ alkanoyl, $C_1$-$C_6$ alkanoyloxy, $C_1$-$C_6$ straight or branched chain alkyl, $C_3$-$C_8$ cycloalkyl, OH, a halogen atom, a straight, branched alkylene chain with 1 to 5 carbon atoms, fluoralkoxy, hydrogen, OR', R' is a halogen atom, $-OR_{23}$ or

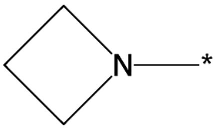,

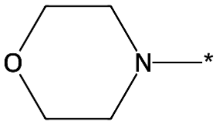,

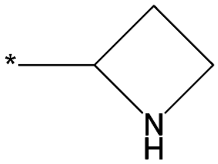,

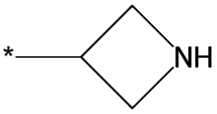,

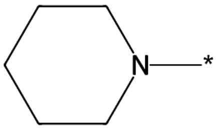,

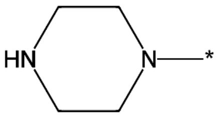,

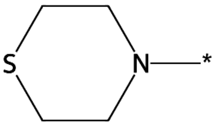,

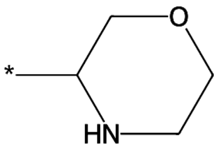,

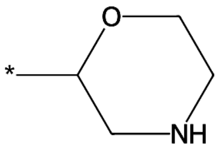,

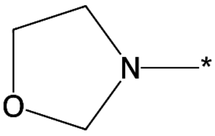,

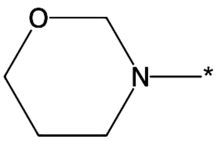 or

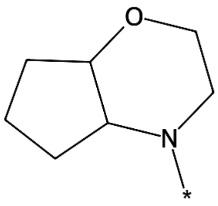;

wherein $R_{23}$ is $C_3$-$C_8$ alkenyl or

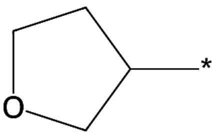,

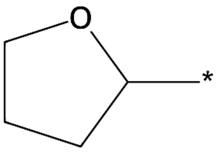,

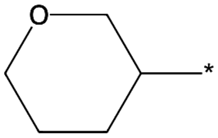,

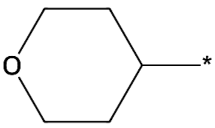,

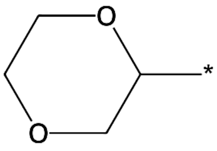,

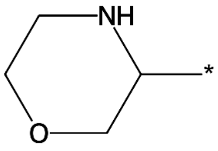,

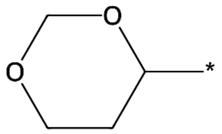

,

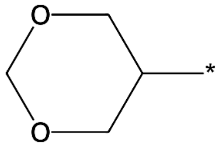

or

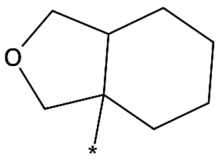

;

$R_{19}$ is $-(CH_2)_2CN$, $-(CH_2)_2N(CH_3)_2$, $-(CH_2)_2O(CH_2)_2OCH_3$, $-(CH_2)_2OCH_3$, $-(CH_2)_3N(CH_3)_2$, $-(CH_2)_3OCH_3$, -BHCN, -CH(Br)CN, $-CH(CH_3)(CH_2)_2OCH_3$, $-CH(CH_3)CH_2N(CH_3)_2$, -CH(CN)OH, $-CH(COOCH_3)CH_3$, $-CH(F)CH_2N(CH_3)_2$, $-CH(NH_2)(CH_2)_2OCH_3$, $-CH(NH_2)CH_2N(CH_3)_2$, $-CH(OCOCH_3)CH_3$, $-CH(OH)CH_2N(CH_3)_2$, $-CH_2C(=NH)OCH_3$, $-CH_2C(=S)N(CH_3)_2$, $-CH_2CH(CH_3)CH_2OCH_3$, $-CH_2CH(F)CH_2OCH_3$, $-CH_2CH(N(CH_3)_2)CH_3$, $-CH_2CH(NH_2)CH_2OCH_3$, $-CH_2CH(OC(=S)S)CH_3$, $-CH_2CH(OC(=S)SH)CH_3$, $-CH_2CH(OH)CH_2OCH_3$, $-CH_2CH(OH)N(CH_3)_2$, $-CH_2CN$, $-CH_2COCH(CH_3)_2$, $-CH_2COCH_2OCH_3$, $-CH_2COOCH(CH_3)_2$, $-CH_2N(C(=NH)CH_3)CH_3$, $-CH_2N(CHO)CH_3$, $-CH_2N(CH_3)CH(CH_3)_2$, $-CH_2N(CH_3)CH_2CH_3$, $-CH_2N(CH_3)_2$, $-CH_2N(COCH_3)CH_3$, $-CH_2NHCH_2OCH_3$, $-CH_2NHCH_3$, $-CH_2NHN(CH_3)_2$, $-CH_2O(CH_2)_2OCH_3$, $-CH_2OC(=NH)CH_3$, $-CH_2OCHO$, $-CH_2OCH_2CH_3$, $-CH_2OCH_2Cl$, $-CH_2OCH_2OCH_3$, $-CH_2OCH_3$, $-CH_2OCOOCH_3$, $-CH_2OF$, $-CH_2OH$, $-CH_2ON(CH_3)_2$, $-CH_2ONHCH_3$, -CN, $-CO(CH_2)_2OCH_3$, $-COCH(Br)CH_3$, $-COCH(OCH_3)CH_3$, $-COCH(SH)CH_3$, $-COCH_2CH(CH_3)_2$, $-COCH_2N(CH_3)_2$, -COCN, $-CON(CH_3)_2$, $-CONHCH(CH_3)_2$, $-COOCH(CH_3)_2$, $-COOCH(Cl)CH_3$, $-COOCH(F)CH_3$, -COOCH(F)F, $-COOCH_2CH_3$, $-COOCH_3$, $-COSCH(CH_3)_2$, $-N(CH_3)(CH_2)_2OCH_3$, $-N(CH_3)CH_2OCH_3$, $-N(CH_3)_2$, $-N(CN)CH_3$, -NCN, $-NHCH_2N(CH_3)_2$, $-NHCH_2OCH_3$, -NHCN, $-O(CH_2)_2CH(CH_3)_2$, $-O(CH_2)_2CH(OH)CH_3$, $-O(CH_2)_2N(CH_3)_2$, $-O(CH_2)_2NHCH_3$, $-O(CH_2)_2OCH_3$, $-O(CH_2)_2OCN$, $-O(CH_2)_2ONH_2$, $-O(CH_2)_2S(CH_3)_2$, $-O(CH_2)_2S(O)CH_3$, $-O(CH_2)_2SCH_3$, $-O(CH_2)_3CH_3$, $-OCH_2N(CH_3)_2$, $-OCH_3$, -OCN, $-OCOCH(CH_3)_2$, $-O_{13}CH_3$, $-SCH_2N(CH_3)_2$, $-SCH_2OCH_3$, -SCN, $=C(CN)NH_2$, =C(CN)OH, $=CH(CH_2)_2OCH_3$, =CHCN, COO-alkyl, COOH, CHO, OH, $C_1$-$C_4$ alkoxy partially or totally substituted with halogen atoms, $C_1$-$C_4$ alkyl partially or totally substituted with halogen atoms, $C_1$-$C_6$ alkanoyl, $C_1$-$C_6$ alkanoyloxy, $C_1$-$C_6$ alkylthio, $C_1$-$C_6$ straight or branched chain alkyl, $C_1$-$C_8$ alkylsulfonyl, $C_3$-$C_8$ alkenyl, $C_3$-$C_8$ cycloalkyl, $C_3$-$C_8$ hydroxyalkyl, $C_3$-$C_8$ straight or branched chain alkyl, OH, a halogen atom, a straight, branched alkylene chain with 1 to 5 carbon atoms, alkynyloxy, carbamoyl, dialkylamino, dimethylamine, dimethylformamide(DMF), fluoralkoxy, hydrogen, substituted/unsubstituded aminos, trihalomethoxy, $C_1$-$C_8$ aminoalkyl, $C_1$-$C_8$ alkylamino, OR', R' is a halogen atom, $-OR_{24}$ or

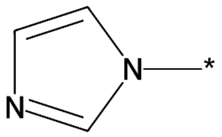

,

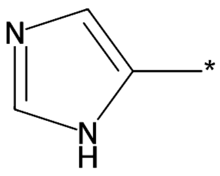

,

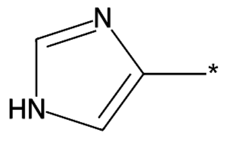

,

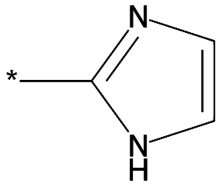
,
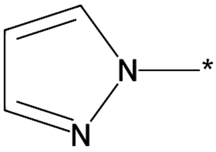
,
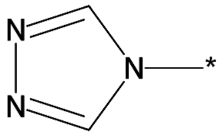
,

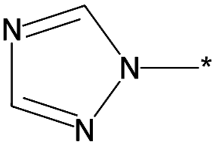
,
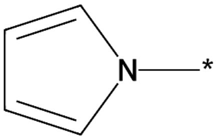
,
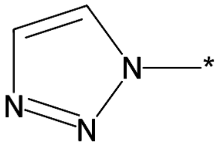
or

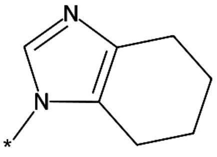
;

wherein $R_{24}$ is $C_3$-$C_8$ alkenyl or

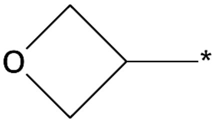
,
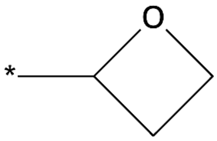
,
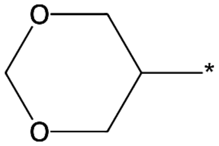
,

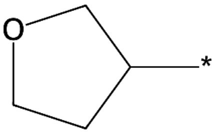
,
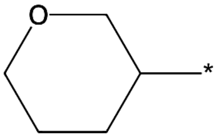
,
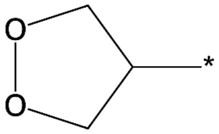
,

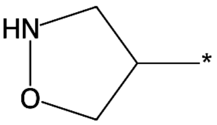
or
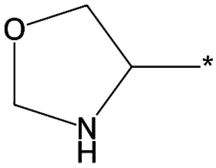
;

$R_{20}$ is -$N(CH_3)_2$, -$NHSO_2CH_3$, $C_1$-$C_4$ alkoxy partially or totally substituted with halogen atoms, $C_1$-$C_4$ alkyl partially or totally substituted with halogen atoms, $C_1$-$C_4$ alkylamino, $C_1$-$C_6$ alkanoyl, $C_1$-$C_6$ alkanoyloxy, $C_1$-$C_6$ straight or branched chain alkyl, $C_3$-$C_8$ cycloalkyl, $NO_2$, OH, a straight or branched alkylene chain with 1 to 5 carbon atoms, alkanesulphonyl, hydrogen or OR', R' is a halogen atom;

$R_5$ is $-N(CH_3)_2$, $C_1$-$C_4$ alkyl partially or totally substituted with halogen atoms, $C_1$-$C_4$ alkylamino, $C_1$-$C_6$ alkanoyl, $C_1$-$C_6$ straight or branched chain alkyl, $C_3$-$C_8$ cycloalkyl, a straight or branched alkylene chain with 1 to 5 carbon atoms or hydrogen;

$X_2$ represents H, C, N, S or O;

$X_1$, $X_3$ and $X_4$ are the same or different and each independently represents C or N.

**Document 4: US 6 248 771 B2**

Claims

1. A compound of formula (I):

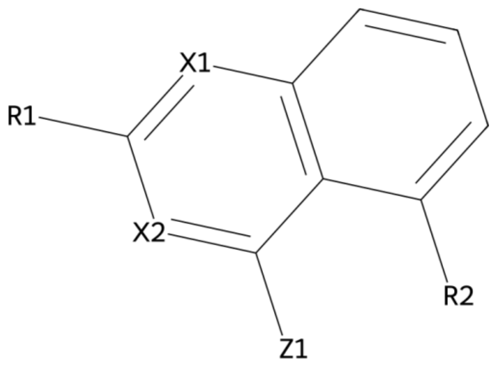

(I)

wherein:

$R_1$ is $C_1$-$C_4$ aminoalkyl, carbamoyl, hydrogen or substituted/unsubstituded aminos;

$R_2$ is 1-pyrrolyl, $C_1$-$C_{12}$ aralkyl, arlthios, benzene ring, hydrogen, phenoxy or phenylamino, -$OR_3$;

wherein $R_3$ is $C_1$-$C_{12}$ aralkyl, $C_1$-$C_6$ straight or branched chain alkyl, Pyrimidine, Pyrrole, Thiazole, benzene ring, cycloalkyl or phenyl optionally substituted with halogen, hydrogen, pyridylmethyl or

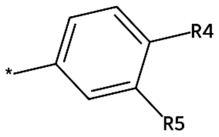

or

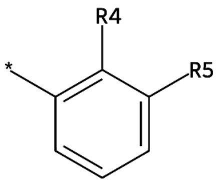

;

wherein $R_4$ is $C_1$-$C_4$ alkoxy partially or totally substituted with halogen atoms, $C_1$-$C_4$ alkyl partially or totally substituted with halogen atoms, $C_1$-$C_6$ alkanoyloxy, OH, hydrogen, trihalomethoxy or OR', R' is a halogen atom;

$R_5$ is $-N(CH_3)_2$, $C_1$-$C_4$ alkoxy partially or totally substituted with halogen atoms, $C_1$-$C_4$ alkyl partially or totally substituted with halogen atoms, $C_1$-$C_6$ alkanoyloxy, $C_1$-$C_6$ alkylthio, $C_1$-$C_6$ straight or branched chain alkyl, $C_3$-$C_8$ hydroxyalkyl, $C_3$-$C_8$ straight or branched chain alkyl, OH, dialkylamino, dimethylamine, dimethylformamide(DMF), hydrogen, trihalomethoxy, $C_1$-$C_8$ aminoalkyl, $C_1$-$C_8$ alkylamino or OR', R' is a halogen atom;

$X_1$, $X_2$ and $Z_1$ are the same or different and each independently represents C or N.
